# Using natural language processing and machine learning to classify health literacy from secure messages: The ECLIPPSE study

**DOI:** 10.1101/406876

**Authors:** Renu Balyan, Scott A. Crossley, William Brown, Andrew J. Karter, Danielle S. McNamara, Jennifer Y. Liu, Courtney R. Lyles, Dean Schillinger

## Abstract

Limited health literacy can be a barrier to healthcare delivery, but widespread classification of patient health literacy is challenging. We applied natural language processing and machine learning on a large sample of 283,216 secure messages sent from 6,941 patients to their clinicians for this study to develop and validate literacy profiles as indicators of patients’ health literacy. All patients were participants in Kaiser Permanente Northern California’s DISTANCE Study. We created three literacy profiles, comparing performance of each literacy profile against a gold standard of patient self-report. We also analyzed associations between the literacy profiles and patient demographics, health outcomes and healthcare utilization. T-tests were used for numeric data such as A1C, Charlson comorbidity index and healthcare utilization rates, and chi-square tests for categorical data such as sex, race, continuous medication gaps and severe hypoglycemia. Literacy profiles varied in their test characteristics, with C-statistics ranging from 0.61-0.74. Relationships between literacy profiles and health outcomes revealed patterns consistent with previous health literacy research: patients identified via literacy profiles as having limited health literacy were older and more likely minority; had poorer medication adherence and glycemic control; and higher rates of hypoglycemia, comorbidities and healthcare utilization. This research represents the first successful attempt to use natural language processing and machine learning to measure health literacy. Literacy profiles offer an automated and economical way to identify patients with limited health literacy and a greater vulnerability to poor health outcomes.

## BACKGROUND AND SIGNIFICANCE

An estimated 30.3 million people in the U.S. had diabetes mellitus (DM) in 2015 according to the Centers for Disease Control and Prevention (2017). Like most chronic conditions, DM self-management can be complex and requires that patients frequently communicate with healthcare providers. Health literacy (HL) is generally defined as a patient’s ability to obtain, process, comprehend and communicate basic health information [1, 2]. DM patients with limited HL have a higher risk of poor health outcomes, including worse blood sugar control, higher complication rates [3], and a greater incidence of hypoglycemia [4, 5]. Poor communication and sub-optimal adherence to medication may explain some of these disparities [6, 7]. Limited HL contributes to preventable suffering, more rapid decline in physical function [8] and related healthcare costs. Online patient portals embedded within electronic health records (EHRs) are now being used widely to bridge in-person encounters and providing support between visits by allowing patients and providers to communicate via secure messages (SMs). The reach and effectiveness of online communication is likely heavily affected by patients’ HL. Limited HL is a barrier to use of patient portals and impacts patients’ evaluation of online health information [9]. However, no research has harnessed SMs to identify patients with limited HL to provide disease management support. Developing scalable tools to identify limited HL without primary data collection burden would be an efficient way to target tailored provider communication and related interventions. The goal of the ECLIPPSE study (Employing Computational Linguistics to Improve Patient-Provider Secure Email exchanges) is to develop patient literacy profiles (LPs) using natural language processing (NLP) tools and machine learning (ML) to classify HL (limited vs. not) in a large sample of SMs from diabetes patients and to assess whether LPs are associated with patient demographics and health outcomes. We hypothesize that patients’ language constructs in portal communications can be harnessed to estimate their health literacy (HL).

Prior research in medical domains has benefitted from the use of NLP and ML, such as representation of clinical narratives, assessing medical articles’ text quality, and developing semantic lexicons for medical language processing [10-17]. Some of the commonly used NLP tools and techniques these studies employed are Apache clinical text analysis and knowledge extraction system (cTAKES) [18], the clinical language annotation, modeling, and processing tool (CLAMP) [19], the medical language extraction and encoding system (MedLee) [20], the Kawasaki disease-NLP (KD-NLP) [21] tool, Flesch-Kincaid Grade level (FKGL) [22], SMOG [23-24] and suitability assessment of materials (SAM) [25] tool. Most of these tools have been used for clinical analyses and not HL; very few ones like Flesch-Kincaid and SMOG use surface-level features centered on relatively shallow lexical and sentential indices.

Despite the increasing use of NLP and ML techniques in health domains, to our knowledge, no study has utilized these techniques to estimate the HL of patients. Kim and Xie [26] carried out a literature survey to identify online health services used by people with limited HL. These authors concluded that there is a need for new HL screening tools. Healthcare delivery systems are recognizing the importance of identifying the subset of patients who have limited HL. Measuring HL, however, requires the use of interviews or questionnaires, rendering the process challenging, especially for larger patient populations. An automated LP based on NLP would provide a more efficient means to identify patients with limited HL. We set out to develop an automated LP prototype that can (a) identify patients with potential HL limitations in an automated way, and (b) determine whether the measures are associated with health outcomes, and (c) deliver feedback to clinicians about the HL skills of patients so that clinicians can modify their language to make SMs to the patients more readable and actionable, thereby improving communication. To accomplish the first two objectives, the current study examines the extent to which patients’ self-reported HL can be estimated using LP models created using NLP and ML techniques.

## MATERIALS AND METHODS

KP has a well-developed and mature patient portal, kp.org. Previous research suggests that patients who access such portals are more likely to have better (a) healthcare utilization [27], (b) medication adherence [28-29] and (c) glycemic (blood sugar) control [30-31]. Among DM patients, better ratings of physician communication are associated with greater SM usage [32]. Limited HL poses a barrier to portal and SM use. However, these disparities are rapidly narrowing. In 2014, 68% of KP DM patients with limited HL and 84% with adequate HL accessed the portal. Overall, 46% used SM in 2014, compared to 30% in 2009. Those with limited HL are rapidly gaining ground, showing a 65% increase in 5 years period compared to a 41% increase for adequate HL (unpublished data). The greatest gains have been among Latinos and African Americans.

### Data source and participants

Data for this study were extracted from the KPNC Diabetes Registry (N∼320,000, as of 01/01/2017). Our sampling frame includes >1 million SMs generated by >150,000 ethnically diverse DM patients and >9,000 clinicians from an integrated delivery system - KPNC. We identified the subset of these patients who completed a survey entitled the Diabetes Study of Northern California (DISTANCE), including providing self-reported HL (N=14,357) [33-35]. DISTANCE involved a survey of DM patients receiving care from KPNC, oversampling minority sub-groups to assess the role of socio-demographic factors on quality of care. The variables in DISTANCE were collected from questionnaires completed via telephone, on-line, or paper and pencil (62% response rate).

We extracted all the SMs (N=1,050,577) exchanged between a patient and all clinicians from KP’s patient portal between 01/01/2006 and 12/31/2015. We then identified those SMs that a patient sent to his or her primary care physician(s). We also removed all patients whose SM lacked sufficient words (<50 words) to provide linguistic coverage and patients who did not have matching DISTANCE survey data. We then removed all SMs written in a language other than English and all SMs identified as written by proxies (i.e., SMs written for the patient by caregivers) [36]. The final cleaned data consisted of 6,941 patients and 283,216 SMs. These SMs were collated into a single file from which we extracted the linguistic features for each patient, aggregating their SMs. These linguistic features were used to predict HL based on self-reported HL scores obtained from survey data.

### Variables

#### Primary predictors: The Linguistic Features

We used a set of 185 linguistic features, derived from the patients’ SMs sent to their clinicians, to predict patients’ self-reported HL and create the LPs. We used NLP tools to select features that measure different language aspects, such as text level information (e.g., number of words in the text, token type ratio), lexical sophistication, syntactic complexity, text cohesion (e.g., connectives, word overlap), and affect (S1 Table). These linguistic aspects have previously been shown to predict literacy levels in non-clinical corpora [37-38]. NLP tools used to extract these features included the Tool for the Automatic Assessment of Lexical Sophistication (TAALES) [39-40], the Tool for the Automatic Analysis of Cohesion (TAACO) [41], the Tool for the Automatic Assessment of Syntactic Sophistication and Complexity (TAASSC) [42-43], the SEntiment ANalysis and Cognition Engine (séance) [44], and the Writing Assessment Tool (WAT) [45-46].

#### Dependent Variable(s): Self-Reported Health Literacy

As a gold standard, we used combinations of self-reported HL items from DISTANCE survey to compute three dependent variable versions of predicted self-reported HL. The survey included the following HL measures: self-reported confidence in filling out medical forms (HLCONF), problems in understanding written medical information (HLPROB), frequency of needing help in reading and understanding health materials (HLHELP); and an original item: problems understanding prescription labels (HLLABELS) (S2 Table). The first three items have been previously validated [47]. Patient responses were collected using a 5-point Likert scale in which a response of 1 referred to “Always” and a response of 5 to “Never.” For our analyses, we combined these items to create different self-reported variables to be able to compare the performance of the linguistic features against different computations of self-reported HL (i.e., combined HL [HLCOMB], trinary summed HL [HLSUMTri], and average HL [HLAVG]; see S2 Table for definitions and computation of these variables).

HLCOMB considers binary forms of three self-reported HL measures (HLPROB2, HLCONF2, and HLHELP2 in Appendix 2); a ‘zero’ score indicates that a patient reports no HL limitations and a ‘one’ that a patient reports limited HL on any one of the three items. HLSUMTri is a trinary variable computed by summing the Likert scale values obtained for HLPROB, HLCONF, and HLHELP. The HLSUMTri variable had three possible values ranging between 0 and 2. Zero (0) indicates a patient with limited HL, whereas one (1) and two (2) represent a patient with marginal and adequate HL, respectively. The HLAVG scores were computed by taking the mean of HLPROB, HLHELP, HLCONF, and HLLABELS (S2 Table).

#### Dependent Variable(s): Health Outcomes

Using data derived from the EHR, we examined medication adherence based on continuous medication gaps (CMG) [48-49], a validated adherence measure of percent time with insufficient medication supply; hypoglycemia (a side effect of DM treatment that can be a marker for poor communication); Hemoglobin A1c (an integrated measure of blood sugar control); and Charlson index [50-51] (a measure of comorbidity and illness severity). Comorbid illness was measured with the Deyo version of the Charlson comorbidity index [52]. We considered patients to have poor adherence if CMG>20% and adequate adherence when CMG≤20% [53]. A1c was based on the most recent value collected after the first SM sent since DISTANCE survey completion, and CMG, severe hypoglycemia and Charlson index were measured the year before the first SM was sent. The occurrence of any hypoglycemia-related ED visit or hospitalization was based on a validated algorithm [54] (any of the following ICD-9 codes: 251.0, 251.1, 251.2, 962.3, or 250.8, without concurrent 259.8, 272.7, 681.XX, 682.XX, 686.9X, 707.1-707.9, 709.3 730.0-730.2, or 731.8 codes). Another set of analysis was conducted for health service utilization, using outpatient clinic visits, emergency room encounters and hospitalizations.

### Statistical analysis

Analyses were conducted to develop three LPs using several supervised ML algorithms [55-59]. We examined links between three summed self-reported HL variables (HLCOMB, HLSUMTri, and HLAVG) and the 185 linguistic predictor variables extracted using the linguistic tools. To perform binary classification, we categorized the summed self-reported HL scores into discrete levels (limited vs. high HL). We trained ML models, including linear discriminant analysis (LDA), support vector machines (SVM), random forests, and artificial neural networks on 70% of the data and tested the model performance on the remaining 30%. In all cases, linguistic features were used to predict the discrete HL levels. Several metrics such as accuracy, sensitivity, specificity, positive and negative predictive values (PPV and NPV), and C-statistic (area under the receiver operator characteristic (ROC) curves) were used as measures of model performance using a split sample approach. The resulting LPs were subsequently validated against self-reported HL items previously collected from the patients via in the DISTANCE survey [34], and the HL-sensitive health outcomes obtained from administrative data from the EHR, described above. We discuss the results of the three models that performed the best for each of the dependent variables.

Lastly, to examine whether the ML approaches resulted in patterns similar to those reported in prior literature on self-reported and directly measured HL, we examined bivariate associations between each of the LP models and demographic, health outcome and healthcare utilization variables using a two-sided p-value at the 0.05 level of significance. Categorical variables such as sex, race, continuous medication gaps [53] and severe hypoglycemia were analyzed using chi-square analysis. Mean comparisons were conducted using t-tests for A1c, Charlson (comorbidity) index [50], healthcare utilization rates.

## RESULTS

### Aggregated Health Literacy Measures

The first analysis to create an LP modeled HLCOMB as the dependent variable. The data for HLCOMB were distributed uniformly, with 3,229 patients having high HL (or no HL limitations), and 3,712 limited HL. The LDA model performed the best for this version of the LP, achieving an accuracy of 60.55% and a C-statistic of 0.63 for the test data (Table 1; bold entries indicate the highest value for a given metric within an LP).

**Table 1:** Classification metric statistics of models for different self-reported Literacy Profiles (Positive class: High HL)

| ML Algorithm for Literacy Profiles | Literacy Profile (Dependent Variable) | Accuracy | C-statistic | Sensitivity | Specificity | Positive Predictive Value (PPV) | Negative Predictive Value (NPV) | # of Predicted limited vs high HL* |
| --- | --- | --- | --- | --- | --- | --- | --- | --- |
| LDA | HLCOMB | 60.55 | 0.63 | 56.10 | 64.42 | 57.83 | 62.78 | 1142 / 939 |
| LDA | HLSUMTri | **63.58** | 0.61 | 39.32 | **79.32** | 55.23 | **66.82** | 1498 / 583 |
| SVM | HLAVG | 62.52 | **0.74** | **75.49** | 47.11 | **62.91** | 61.79 | 725 / 1356 |

\* The numbers are a function of sample size for test set only

The second analysis considered HLSUMTri as the dependent variable to create an LP. Since the HLSUMTri variable had three possible values (classes), we used multiclass classification. The accuracy of the models was lower and ranged between 50.67% and 54.23%. SVM achieved the highest accuracy. However, SVM classified all the instances as marginal or adequate HL. To determine how these algorithms performed using binary classification, we combined the inadequate (0) and marginal (1) HL instances and re-classified these as limited (0+1) HL, while the high (2) HL cases were retained. In binary classification, the LDA model performed the best, and the results for these models were better than the multiclass classification results. The LDA model achieved an accuracy of 63.58% and a C-statistic of 0.61. However, the C-statistic was lower than the LDA model of the LP trained using HLCOMB, as was its sensitivity (39.32% vs. 56.10%, Table 1).

For the third analysis, we considered the HLAVG scores as the dependent variable to create an LP. The data set included 3,173 limited HL and 3,768 high HL instances. Accuracy and other metrics were observed for the SVM version of this LP: accuracy and c-statistic for SVM model were 62.52% and 0.74 respectively. While the specificity was lower, it achieved the greatest balance in PPV and NPV (Table 1).

### Linguistic Characteristics

The LP models generally showed that patients with predicted limited HL produced messages having fewer words, and those words were less sophisticated (i.e., more concrete) and demonstrated less lexical diversity (i.e., greater repetition of words). Additionally, patients with limited predicted HL produced more words that expressed negative affect (i.e., more words related to failure and fewer positive words). Lastly, limited predicted HL patients focused less on personal language, using a greater incidence of third person pronouns and fewer first person pronouns.

### Demographics

The average age of our study population at the time of the DISTANCE study was 56.8 (±10); 54.3% were male and 32.2% were white. When applying the ML model-derived LPs to the validation dataset, we found patterns that matched previously observed relationships between patient characteristics and HL. For example, patients identified by the LPs to have limited HL were 1-3 years older than high HL patients. In addition, 70.8-76.1% of the predicted limited HL patients were non-white, compared to 59.9-63.5% of high HL patients (Table 2), and 84.7-88.7% of patients with predicted limited HL had high school diplomas compared to 93.4-95% of patients with high HL.

**Table 2:** Demographics (Sex %, Race % and Age – Mean (SD))

| ML Algorithm for Literacy Profiles | Literacy Profile (Dependent Variable) | Sex - Men % |  |  | Race – White % |  | Age at Survey – Mean (SD) |  | P-value |
| --- | --- | --- | --- | --- | --- | --- | --- | --- | --- |
|  |  | Limited HL | High HL | P-value | Limited HL | High HL | Limited HL | High HL |  |
| LDA | HLCOMB | 54.9 | 53.7 | 0.32 | 25.5 | 40.0 | 57.91 (10.0) | 55.53 (9.66) | < 0.001 |
| LDA | HLSUMTri | 55.8 | 53.6 | 0.08 | 29.2 | 40.1 | 57.34 (10.0) | 55.43 (9.50) | < 0.001 |
| SVM | HLAVG | 53.6 | 56.2 | 0.06 | 23.9 | 36.5 | 58.88 (9.98) | 55.74 (9.74) | < 0.001 |

### Health Outcomes

To evaluate whether the LP scores were associated with health outcomes in the anticipated directions, we linked these modeled LP scores to outcomes previously found to be associated with measured HL. The results for medication adherence for LP models using HLCOMB and HLSUMTri lacked significance, whereas the model for HLAVG was statistically significant (Table 3). Patients with limited HL based on this LP were more likely to have poor medication adherence than high HL patients (24.5%-25.6% vs. 23.2%-23.4%). Patients predicted to have limited HL had higher severe hypoglycemia rates in all the models, with SVM distinguishing the most. In sum, the SVM version of the LP HLAVG appeared to be the LP that performed best.

**Table 3:** Poor adherence and Hypoglycemia (%)

| ML Algorithm for Literacy Profiles | Literacy Profile (Dependent Variable) | Poor medication adherence (%) |  |  | Severe Hypoglycemia (%) |  |  |
| --- | --- | --- | --- | --- | --- | --- | --- |
|  |  | Limited HL | High HL | P-value | Limited HL | High HL | P-value |
| LDA | HLCOMB | 24.9 | 23.3 | 0.143 | 4.0 | 2.0 | < 0.001 |
| LDA | HLSUMTri | 24.5 | 23.2 | 0.296 | 3.5 | 2.1 | < 0.001 |
| SVM | HLAVG | 25.6 | 23.4 | 0.047 | 5.1 | 2.0 | < 0.001 |

Table 4 shows that patients predicted to have limited HL as measured by the LP HLAVG had poorer glycemic control. Patients with predicted limited HL had higher prevalence of comorbid conditions compared to those with high HL (up to 0.63 more on Charlson score). Again, the SVM version of the LP HLAVG appeared to be the LP that performed best.

**Table 4:** A1c and Charlson Index - Mean (SD)

| ML Algorithm for Literacy Profiles | Literacy Profile (Dependent Variable) | A1c |  |  | Charlson Index |  |  |
| --- | --- | --- | --- | --- | --- | --- | --- |
|  |  | Limited HL | High HL | P-value | Limited HL | High HL | P-value |
| LDA | HLCOMB | 7.51 (1.56) | 7.48 (1.50) | 0.371 | 2.44 (1.78) | 1.99 (1.39) | < 0.001 |
| LDA | HLSUMTri | 7.50 (1.54) | 7.49 (1.52) | 0.786 | 2.34 (1.71) | 1.94 (1.34) | < 0.001 |
| SVM | HLAGV | 7.55 (1.57) | 7.47 (1.51) | 0.038 | 2.65 (1.91) | 2.02 (1.41) | < 0.001 |

### Healthcare Service Utilization

Finally, analyses of healthcare service utilization rates demonstrated that patients with predicted limited HL had on average 10 outpatient clinic visits annually, compared to an average of 8 to 9 among patients with high HL. Similar differences were found for emergency room visits (0.53 vs 0.31) and inpatient hospitalizations (0.25 vs 0.13; see Table 5). These were significant for all models, although the differences in emergency room visits and inpatient hospitalizations were most robust for the SVM HLAVG version.

**Table 5:** Healthcare Service Utilization (outpatient clinic visit, emergency room encounter and hospitalization – Mean (SD))

| ML Algorithm for Literacy Profiles | Literacy Profile (Dependent Variable) | Outpatient clinic visit |  | ED visits |  | Hospitalization |  | P-value |
| --- | --- | --- | --- | --- | --- | --- | --- | --- |
|  |  | Limited HL | High HL | Limited HL | High HL | Limited HL | High HL |  |
| LDA | HLCOMB | 10.02 (10.4) | 8.76 (8.76) | 0.46 (1.07) | 0.30 (0.75) | 0.21 (0.68) | 0.13 (0.51) | < 0.001 |
| LDA | HLSUMTri | 9.69 (10.0) | 8.79 (8.81) | 0.42 (1.00) | 0.31 (0.75) | 0.19 (0.63) | 0.14 (0.56) | < 0.001 |
| SVM | HLAGV | 10.29 (10.7) | 9.01 (9.16) | 0.53 (1.20) | 0.31 (0.76) | 0.25 (0.73) | 0.13 (0.54) | < 0.001 |

## DISCUSSION

The objective of the study was to examine the extent to which HL can be estimated through the linguistic features of DM patients’ secure messages. We compared three LPs modeled from different derivations of patients’ self-reported HL using multiple ML algorithms to determine the LP that best predicted self-reported HL. The SVM LP model for HLAVG performed quite well for all the metrics except specificity, and it generated the best balance with respect to PPV and NPV. In addition, HLAVG predicted that about 1/3 of patients have limited HL, consistent with prior research. With respect to confirmation of previous correlations between accepted measures of HL and health outcomes, the LP derived from the HLAVG SVM model also performed the best.

Overall, we found that several linguistic features that measure different language aspects of SMs derived from electronic patient portals yielded models that predicted self-reported HL with a modest but acceptable degree of accuracy. Together, these features, including less sophisticated and less positive language, provide us with a language profile of limited HL patients. While the linguistic features we included have been previously studied to classify literacy [37-38], the texts that have been assessed have not been derived from e-mail messages. We found that combinations of language features can be applied to SMs to successfully distinguish patients based on self-reported metrics of HL. To our knowledge, this represents the first successful attempt to use NLP to identify patients who have higher likelihoods of self-reported limited HL and vulnerability to worse health outcomes.

The ultimate goal of this work is to develop tools to improve communication between clinicians and patients so as to foster “shared meaning”. Measuring HL has traditionally been extremely challenging at both the individual and population levels, given the time and personnel demands intrinsic to current HL measurement approaches. An automated LP could provide an efficient means to help identify the subpopulation of patients with limited HL. Given that limited HL is an important and potentially remediable factor influencing the incidence of, complication rates of, and mortality from DM and other chronic diseases, developing a valid method for rapid HL assessment likely represents a significant accomplishment with potentially broad public health and clinical benefits. For instance, identifying patients with potentially limited HL could prove useful for alerting physicians about potential difficulties in comprehending written instructions. This lack of comprehension is particularly critical when there are significant drug safety concerns, e.g., anticoagulants and insulin. Additionally, patients identified as having limited HL could be flagged to receive follow up communications to ensure understanding of medication instructions and adherence.

### Limitations

Our study has important limitations. While our patient sample was large and ethnically diverse, and we studied a large number of patients’ SMs, we were only able to analyze those patients who had engaged in SM with their clinicians. As such, the SM-based method used in this study can only be applied to patients who use SM. However, recent data suggest that patients with limited HL are accelerating in their use of patient portals, and over 2/3 of Kaiser diabetes patients with limited HL now use the patient portal. In addition, we limited the study to only English SMs, excluded second language patients who may have limited HL. At the time of this study, KPNC did not have a Spanish language portal.

Our future research will compare performance of these LP models with novel LPs derived from (a) linguistic expert ratings of SMs and (b) existing linguistic indices that estimate literacy; we will examine the relative performance of these LPs in safety net healthcare systems, as well as in patient populations with conditions other than DM. In addition, while limited HL is more heavily concentrated in safety net healthcare settings, this phase of our research involved a fully insured population (Kaiser Permanente) because of the availability of extensive linguistic and health-related data. However, Kaiser Permanente has a sizable Medicaid population, and over 1/4 of their patients have limited HL [4, 47]. Moreover, Kaiser Permanente members are ethnically diverse and largely representative of the U.S. population, with the exception of extremes of income. Finally, while our cross-sectional bivariate analyses with respect to health outcomes were confirmatory, future work will utilize longitudinal data to examine the extent to which LPs may be causally associated with changes in health.

## CONCLUSION

Population management is increasingly incorporating predictive models and derived scores as a means of risk stratifying and targeting care. Our LPs offer healthcare delivery systems a novel, automated, and economical way to identify the subset of patients who have higher likelihoods of having limited HL. Based on our results, we recommend that researchers and health system planners interested in using NLP to identify limited HL use the version of the LP that we have named SVM HLAVG. The LP-derived information could be used to tailor and target both communication and clinical interventions at the health system level. In addition, LPs could be employed as a provider alert in the EHR to improve individual-level communication, or could be harnessed to provide automated feedback to clinicians as they are composing SMs. Insofar as the subset of patients using SM is large and rapidly growing, a literacy profile will soon be calculable on the majority of patients. While the LP is only a proxy for actual barriers to health-related communication, our research demonstrates that LPs are modestly associated with both self-reported HL as well as health outcomes previously shown to be sensitive to HL (e.g., medication adherence, Hemoglobin A1c, hypoglycemia, comorbidities, and utilization). Our future work will (1) compare alternative methods to estimate HL, including those derived from expert ratings and previously validated linguistic indices, (2) develop similar measures for clinicians’ SMs to measure linguistic discordance with patients, (3) determine if automated feedback to clinicians improves SM linguistic concordance, and (4) extend this research to safety net healthcare settings.

## CONTRIBUTORS

JL, RB and SC collected and cleaned the data. AK, DS, JL, RB and SC analyzed and interpreted the data. AK, CL, DM, DS, RB, and SC contributed to the study design. AK, CL, DM, DS, JL, RB, SC and WB drafted the report. AK, DM, DS and WB provided critical revision of the article. All authors gave input to the final version and provided final approval of the version to be published.

## REFERENCES

1 Grossman EG, Office of the Legislative Counsel. Patient Protection and Affordable Care Act, Edited by U.D.o.H.H. Services, Department of Health & Human Services, Washington, DC, USA, 2010.

2 Schillinger D, McNamara DS, Crossley SA, et al. The Next Frontier in Communication and the ECLIPPSE Study: Bridging the Linguistic Divide in Secure Messaging. Journal of Diabetes Research, Vol. 2017, Article ID 1348242, 9 pages. doi: 10.1155/2017/1348242.

3 Schillinger D, Grumbach K, Piette J, et al. Association of health literacy with diabetes outcomes. Jama. 2002 Jul 24;288(4):475–82.

4 Sarkar U, Karter AJ, Liu JY, et al. Hypoglycemia is more common among type 2 diabetes patients with limited health literacy: the Diabetes Study of Northern California (DISTANCE). Journal of general internal medicine. 2010 Sep 1;25(9):962–8.

5 Schillinger D, Bindman A, Wang F, et al. Functional health literacy and the quality of physician–patient communication among diabetes patients. Patient education and counseling. 2004 Mar 1;52(3):315–23.

6 Bailey SC, Brega AG, Crutchfield TM, et al. Update on health literacy and diabetes. The Diabetes Educator. 2014 Sep;40(5):581–604.

7 Bauer AM, Schillinger D, Parker MM, et al. Health literacy and antidepressant medication adherence among adults with diabetes: the diabetes study of Northern California (DISTANCE). Journal of general internal medicine. 2013 Sep 1;28(9):1181–7.

8 Smith SG, O’conor R, Curtis LM, et al. Low health literacy predicts decline in physical function among older adults: findings from the LitCog cohort study. J Epidemiol Community Health. 2015 Jan 8:jech–2014.

9 Diviani N, van den Putte B, Giani S, et al. Low health literacy and evaluation of online health information: a systematic review of the literature. Journal of medical Internet research. 2015 May;17(5).

10 Carrell DS, Cronkite D, Palmer RE, et al. Using natural language processing to identify problem usage of prescription opioids. International journal of medical informatics. 2015 Dec 1;84(12):1057–64.

11 Demner-Fushman D, Chapman WW, McDonald CJ. What can natural language processing do for clinical decision support?. Journal of biomedical informatics. 2009 Oct 1;42(5):760–72.

12 Friedman C, Johnson SB, Forman B, et al. Architectural requirements for a multipurpose natural language processor in the clinical environment. In Proceedings of the Annual Symposium on Computer Application in Medical Care 1995 (p. 347). American Medical Informatics Association.

13 Heintzelman NH, Taylor RJ, Simonsen L, et al. Longitudinal analysis of pain in patients with metastatic prostate cancer using natural language processing of medical record text. Journal of the American Medical Informatics Association. 2012 Nov 9;20(5):898–905.

14 Johnson SB. A semantic lexicon for medical language processing. Journal of the American Medical Informatics Association. 1999 May 1;6(3):205–18.

15 Nadkarni PM, Ohno-Machado L, Chapman WW. Natural language processing: an introduction. Journal of the American Medical Informatics Association. 2011 Sep 1;18(5):544–51.

16 Osborne JD, Wyatt M, Westfall AO, et al. Efficient identification of nationally mandated reportable cancer cases using natural language processing and machine learning. Journal of the American Medical Informatics Association. 2016 Mar 28;23(6):1077–84.

17 Strauss JA, Chao CR, Kwan ML, et al. Identifying primary and recurrent cancers using a SAS-based natural language processing algorithm. Journal of the American Medical Informatics Association. 2012 Aug 2;20(2):349–55.

18 Savova GK, Masanz JJ, Ogren PV, et al. Mayo clinical Text Analysis and Knowledge Extraction System (cTAKES): architecture, component evaluation and applications. Journal of the American Medical Informatics Association. 2010 Sep 1;17(5):507–13.

19 Soysal E, Wang J, Jiang M, et al. CLAMP–a toolkit for efficiently building customized clinical natural language processing pipelines. Journal of the American Medical Informatics Association. 2017 Nov 24.

20 Friedman C, Johnson SB, Forman B, et al. Architectural requirements for a multipurpose natural language processor in the clinical environment. In Proceedings of the Annual Symposium on Computer Application in Medical Care 1995 (p. 347). American Medical Informatics Association.

21 Doan S, Maehara CK, Chaparro JD, et al. Building a natural language processing tool to identify patients with high clinical suspicion for Kawasaki disease from emergency department notes. Academic Emergency Medicine. 2016 May;23(5):628–36.

22 Flesch R. A new readability yardstick. Journal of applied psychology. 1948 Jun;32(3):221.

23 Mc Laughlin GH. SMOG grading-a new readability formula. Journal of reading. 1969 May 1;12(8):639–46.

24 Doak LG, Doak CC. Lowering the silent barriers to compliance for patients with low literacy skills. Promoting Health. 1987;8(4):6–8.

25 Doak CC, Doak LG, Root JH. Teaching patients with low literacy skills 2nd ed. Philadelphia, PA: JB Lippincott; 1996.

26 Kim H, Xie B. Health literacy in the eHealth era: a systematic review of the literature. Patient education and counseling. 2017 Jun 1;100(6):1073–82.

27 Reed M, Huang J, Brand R, et al. Implementation of an outpatient electronic health record and emergency department visits, hospitalizations, and office visits among patients with diabetes. Jama. 2013 Sep 11;310(10):1060–5.

28 Lyles CR, Sarkar U, Schillinger D, et al. Refilling medications through an online patient portal: consistent improvements in adherence across racial/ethnic groups. Journal of the American Medical Informatics Association. 2015 Sep 2;23(e1):e28–33.

29 Sarkar U, Lyles CR, Parker MM, et al. Use of the refill function through an online patient portal is associated with improved adherence to statins in an integrated health system. Medical care. 2014 Mar;52(3):194.

30 Harris LT, Koepsell TD, Haneuse SJ, et al. Glycemic control associated with secure patient-provider messaging within a shared electronic medical record: a longitudinal analysis. Diabetes care. 2013 Sep 1;36(9):2726–33.

31 Reed M, Huang J, Graetz I, et al. Outpatient electronic health records and the clinical care and outcomes of patients with diabetes mellitus. Annals of Internal Medicine, 2012 Oct 2, 157(7): 482–9.

32 Lyles CR, Sarkar U, Ralston JD, et al. Patient–provider communication and trust in relation to use of an online patient portal among diabetes patients: the diabetes and aging study. Journal of the American Medical Informatics Association. 2013 May 15;20(6):1128–31.

33 Chew LD, Griffin JM, Partin MR, et al. Validation of screening questions for limited health literacy in a large VA outpatient population. Journal of general internal medicine. 2008 May 1;23(5):561–6.

34 Moffet HH, Adler N, Schillinger D, et al. Cohort Profile: The Diabetes Study of Northern California (DISTANCE)—objectives and design of a survey follow-up study of social health disparities in a managed care population. International journal of epidemiology. 2008 Mar 7;38(1):38–47.

35 Ratanawongsa N, Karter AJ, Parker MM, et al. Communication and medication refill adherence: the Diabetes Study of Northern California. JAMA internal medicine. 2013 Feb 11;173(3):210–8.

36 Wagatha S, Crossley SA, Karter AJ, et al. Caregiving for Patients with Diabetes in the Era of Secure Messaging: Findings from the ECLIPPSE Study. Society of General Internal Medicine Annual Meeting. April 11, 2018. Denver, CO.

37 Crossley SA, Allen LK, Snow EL, et al. Incorporating learning characteristics into automatic essay scoring models: What individual differences and linguistic features tell us about writing quality. Journal of Educational Data Mining. 2016;8(2):1–9.

38 Crossley SA, Allen LK, McNamara DS. A Multi-Dimensional analysis of essay writing. Multi-Dimensional Analysis, 25 years on: A tribute to Douglas Biber. 2014 Jul 15;60:197.

39 Kyle K, Crossley SA. Automatically assessing lexical sophistication: Indices, tools, findings, and application. Tesol Quarterly. 2015 Dec 1;49(4):757–86.

40 Kyle K, Crossley S, Berger C. The tool for the automatic analysis of lexical sophistication (TAALES): version 2.0. Behavior research methods. 2017 Jul 11:1–7.

41 Crossley SA, Kyle K, McNamara DS. The tool for the automatic analysis of text cohesion (TAACO): Automatic assessment of local, global, and text cohesion. Behavior research methods. 2016 Dec 1;48(4):1227–37.

42 Kyle K. Measuring syntactic development in L2 writing: Fine grained indices of syntactic complexity and usage-based indices of syntactic sophistication.

43 Crossley SA, Skalicky S, Dascalu M, et al. Predicting text comprehension, processing, and familiarity in adult readers: new approaches to readability formulas. Discourse Processes. 2017 Jul 4;54(5-6):340–59.

44 Crossley SA, Kyle K, McNamara DS. Sentiment Analysis and Social Cognition Engine (SEANCE): An automatic tool for sentiment, social cognition, and social-order analysis. Behavior research methods. 2017 Jun 1;49(3):803–21.

45 Crossley SA, Roscoe RD, McNamara DS. Using Automatic Scoring Models to Detect Changes in Student Writing in an Intelligent Tutoring System. In FLAIRS Conference 2013 May 19.

46 McNamara DS, Crossley SA, Roscoe R. Natural language processing in an intelligent writing strategy tutoring system. Behavior research methods. 2013 Jun 1;45(2):499–515.

47 Sarkar U, Schillinger D, López A, et al. Validation of self-reported health literacy questions among diverse English and Spanish-speaking populations. Journal of general internal medicine. 2011 Mar 1;26(3):265–71.

48 Steiner JF, Koepsell TD, Fihn SD, et al. A general method of compliance assessment using centralized pharmacy records: description and validation. Medical care. 1988 Aug 1:814–23.

49 Steiner JF, Prochazka AV. The assessment of refill compliance using pharmacy records: methods, validity, and applications. Journal of clinical epidemiology. 1997 Jan 1;50(1):105–16.

50 Charlson ME, Pompei P, Ales KL, et al. A new method of classifying prognostic comorbidity in longitudinal studies: development and validation. Journal of chronic diseases. 1987 Jan 1;40(5):373–83.

51 Charlson M, Szatrowski TP, Peterson J, et al. Validation of a combined comorbidity index. Journal of clinical epidemiology. 1994 Nov 1;47(11):1245–51.

52 Deyo RA, Cherkin DC, Ciol MA. Adapting a clinical comorbidity index for use with ICD-9-CM administrative databases. Journal of clinical epidemiology. 1992 Jun 1;45(6):613–9.

53 Raebel MA, Schmittdiel J, Karter AJ, et al. Standardizing terminology and definitions of medication adherence and persistence in research employing electronic databases. Medical care. 2013 Aug;51(8 0 3):S11.

54 Ginde AA, Blanc PG, Lieberman RM, et al. Validation of ICD-9-CM coding algorithm for improved identification of hypoglycemia visits. BMC endocrine disorders. 2008 Dec;8(1):4.

55 Balyan R, McCarthy KS, McNamara DS. Combining Machine Learning and Natural Language Processing to Assess Literary Text Comprehension. In A. Hershkovitz & L. Paquette (Eds.). In Proceedings of the 10th International Conference on Educational Data Mining (EDM), Wuhan, China: 2017. International Educational Data Mining Society.

56 Han J, Pei J, Kamber M. Data mining: concepts and techniques. Elsevier; 2011 Jun 9

57 Joachims T. Text categorization with support vector machines: Learning with many relevant features. In European conference on machine learning 1998 Apr 21 (pp. 137–142). Springer, Berlin, Heidelberg.

58 Mitchell TM. Machine learning. 1997. Burr Ridge, IL: McGraw Hill. 1997;45(37):870–7.

59 Schölkopf B, Smola AJ. Learning with kernels: support vector machines, regularization, optimization, and beyond. MIT press; 2002.

60 McCarthy PM. An assessment of the range and usefulness of lexical diversity measures and the potential of the measure of textual, lexical diversity (MTLD). Dissertation Abstracts International. 2005;66:12.

61 Malvern D, Richards BJ, Chipere N, et al. Lexical diversity and language development. New York: Palgrave Macmillan; 2004.

